# Translation coupled mRNA-decay is a function of both structural and codon level characteristics

**DOI:** 10.1101/2024.02.28.582446

**Authors:** Sudipto Basu, Suman Hait, Sudip Kundu

## Abstract

The majority of mRNA degradation occurs co-translationally. Several works in the past elucidated the role of codon composition in regulating co-translational mRNA decay. Integration of mRNA sequence, structure and ribosomal density unravels common regulatory factors of translational and degradation and helps in understanding the intricate association between these two important processes. Co-translational degradation is a two-step process, involving translational stalling and mRNA release for degradation. Our findings highlight the role of Codon Adaptation Index, a sequence-level feature that serves as the primary determinant of translation rates facilitating transcript release from translational machinery upon stalling. Concurrently, cellular endonucleases targeting Internal Unstructured Segments, facilitating easy degradation of the stalled mRNA transcripts, influencing their half-lives across the genome and over evolutionary timescales.

## 1. Introduction

Controlled degradation of biomolecules is essential for maintaining cellular metabolism and homeostasis. mRNA homeostasis is compromised in various neurological diseases, through (i) reduced expression levels and (ii) the increased tendency of the damaged mRNA to aggregate [1]. The pre-requisites for cytoplasmic degradation of mRNA are deadenylation [2] followed by decapping[3], which makes them vulnerable to nuclease action by different 5′ - exonucleases [4–6] and 3′ -exonucleases [7,8]. Alternatively, mRNAs can also be degraded by endo-nucleolytic cleavage [9–11]. A previous study has elucidated the presence of terminal and internal unstructured segments facilitates the action of exonuclease and endonuclease respectively [12].

Cytosolic mRNA spends a major period of its lifecycle shuttling between ribosomal bound and unbound states for translation, so a major gateway to redirect the degradation of mature mRNA is through a co-translation surveillance system[13]. The major co-translational surveillances are: NMD (nonsense-mediated decay), NSD (no-stop decay), and NGD (no-go decay). While NMD or NSD are very specific to their targets, NGD is a broad term encompassing various mRNA translational complications. Evidence suggests that the major targets of NGD are transcripts which account for any form of translational perturbations through a specific string of codons or some higher order mRNA structures within the transcripts[14–16]. Slow translation of non-optimal codon (NOC) directs mRNA for codon-optimality mediated mRNA decay (COMD)[16]. Nevertheless, an extended duration of ribosomal arrests due to presence of NOC can result in NGD[16]. Therefore, the mechanisms of COMD and NGD suggest that mRNAs with smooth translation tend to be more stable.

Smooth translation is a result of abundant cellular concentration of aa-tRNA which ensures minimal wait period at a codon for the ribosomes. The landmark study by Sharp and Li[17] has broadly elucidated codon adaptation indices (CAI and tAI) used to estimate the relative codon composition of a mRNA. The dependency of certain codons in elevating mRNA stability has been previously elucidated in different organisms, including *Saccharomyces cerevisiae*[18], *Schizosaccharomyces pombe*[19], *Trypanosoma brucei*[20], *Drosophila melanogaster*[21], *Danio rerio*[22,23], humans[24,25] and in specific cell-lines (Chinese Hampster Ovary cells and Hela)[26]. In all the mentioned studies, a pearson correlation (r) is calculated between the frequency of each codon within a transcript with the corresponding half-lives. The estimated r-value is used to classify codons as stabilizing (r > 0) or destabilizing (r < 0). Also, it has been observed the codons which classify as stabilizing have a higher adaptation index, while the destabilizing codons have a lower adaptation index [18,19,27].

Several pioneer works on different oncogenes including c-myc [28–30], c-fos [31], jun [32] and myb [33] indicated that translational blockade could result in degradation of that mRNA. Later, it has been shown that this mechanism is not only limited to oncogene-related transcripts, rather also observed at genome scale in multiple organisms [18,19,34–36]. Now it is well accepted that a majority of the mRNA decay occurs co-translationally[37]. Moreover, translational regulation and gene expression are regulated at codon level and the codon composition of a mRNA transcripts also has a substantial effect on its stability. However, codon composition couldn’t explain all the variations in mRNA half-lives[38,39]. Rather, the presence of intrinsically unstructured segments and sequestration play an important role in the observed variations [12,40]. Since endonucleases prefer unstructured segments and help in degradation of all translationally perturbed transcripts [41,42], we hypothesized that the translation-coupled mRNA decay will be a function of both structural and codon level characteristics, which has not been examined extensively till date. Here, we integrate (i) mRNA sequence and structural information and (ii) ribosomal density to gain a profound insight on their synergistic effect on both (a) translation rates and (b) mRNA half-lives in yeast. Alongside, this study is crafted to present key concepts of regulation during translation coupled mRNA decay/COMD/NGD. Our results indicate that (i) there are several regulators to control both the translational rates and half-lives of mRNA; (ii) among these regulators, the relative occurrence of optimal codons (reflected in higher/lower values of CAI, a sequence level feature) and the presence of internal unstructured segments (IUS) are major determinants of mRNA translational rates and half-lives, respectively. Results further suggest that mRNAs with lower CAI values have higher probability of early release from ribosome and are then subjected to degradation machinery including cellular endonucleases that identify internal unstructured segments within transcript and govern mRNA half-lives at genome-scale and throughout evolution.

## 2. Materials and Methods

### 2.1 Data collection

#### 2.1.1 Half-lives and translation rates data

Two non-redundant half-lives datasets for *Saccharomyces cerevisiae* (yeast) are used for this study[18,43] (**Data S1**). The data for translation elongation time used for our analysis are taken from [44]. Translation rates is computed as the length of the transcript translated in unit time 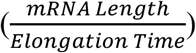, where L indicates the mRNA length and T indicates the elongation time (**Data S1**).

#### 2.1.2 Ribosomal Density

Ribosomal footprinting data provides a quantitative measurement of the number of ribosomes stationed on a mRNA transcript [45]. For the present study, we have obtained the transcript-wise ribosomal profiling reads values and the cytoplasmic RNA abundance data from the GEO database (GEO ID: GSE75897) [46](**Data S1**). The abundance of each transcript is normalized by the ribosomal profiling reads value to obtain the real ribosomal density (RD) for a transcript. There are four sets of ribosomal density data (Unselected, Dynabeads, Ribominus, and Ribozero) based on the different cytoplasmic RNA-seq purification methods. All the primary analysis are performed using Ribozero, since they exhibit a better coverage from 5′ to 3′ -end on the transcriptome[46] (**Data S1**).

### 2.2 Sequence Based-features

#### 2.2.1 Adaptation indices

We chose multiple sequence-based parameters, the values for some of them are taken from literatures, while others are computed in this analysis. The previously features are Codon adaptation index (CAI) and tRNA adaptation index (tAI) [47]. CAI is a classical index where it utilizes a set of reference genes (generally housekeeping genes are preferred) to assess the relative usage of each codon within those genes, and a score is calculated for each of the genes from the usage frequency of each codon within that gene[17]. The generic idea that housekeeping genes possess a higher CAI index, and the codons within them are selected to have a higher translational efficiency (**Data S1**). tAI is also a measure of the translatability of a gene, which considers the intracellular concentration of tRNA molecules and the efficacy of the codon-anticodon pairing[48] (**Data S1**). Hence, a high tAI score for a gene indicates that a codon content for which the corresponding cognate tRNA is more abundant within the cells.

#### 2.2.2 Estimating CSC

Codon stabilization coefficient (CSC) is calculated using the experimental half-lives values, and the annotated yeast transcriptome. For each of the sixty-one codons, the frequencies of a codon (say X) within all the transcripts are mapped with the half-lives of the transcripts. The pearson correlation between the frequencies and the half-lives gives the CSC for that specific codon [18,19,27]. The value of CSC ranges from -1 to +1. In this study, the CSC score is utilized to classify a codon as stabilizing or destabilizing (*p <* 0.001) (**Fig. S1A**)[18,19,27,34]. The initiator codon is always left out for this calculation.

In order to assign each transcript with a unified CSC, we introduced tCSC. tCSC is defined as the summation of all coefficients normalized by the number of codons excluding the initiator and stop codon (**Data S1**). For a transcript with *l* number of codons, tCSC is expressed as:

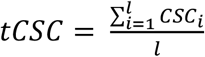

Estimation of tCSC is performed on the CDS region of an mRNA.

### 2.3 Estimating Internal Unstructured Segments

PARS score represents the structuredness of a mRNA at single nucleotide resolution, with the aid of nucleotide digestion and high-throughput sequencing[49]. A positive score (PARS > 0) indicates that the nucleotide site is double-stranded (or base-paired), whereas, a negative score (PARS > 0) indicates that the nucleotide site is single-stranded (or unpaired) conformation. Using PARS data, we have generated an Internal Unstructured Segment (IUS), a derived parameter calculated using a window of 12 nucleotides long moving from 5’ to 3’ end of the transcript with 6 nucleotides slide at a time, similar to the previously established method[12] **(Data S1)**. IUS is calculated as the normalized difference between the number of unstructured and structured segments within a transcript 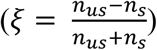 Here, ns and denote the number of unstructured and structured segments in the transcripts. ξ is calculated for different threshold percentages. For example, in a window of 12 nucleotides, if more than 60% of the nucleotides can form base pairs (PARS score > 0), and less than 40% nucleotides are in an unpaired state (PARS score ≤ 0), we denote it as ξ _s60us40_. In a similar way, different percentage threshold is used for counting both structured and unstructured segments and calculating ξ score for each transcript. We found that the ξ value for different structured and unstructured threshold are (i) highly correlated among themselves and (ii) show similar correlation range with mRNA translation rates and half-lives (**Fig. S1B**). Hence, we have performed all the analysis using a representative ξ _s60us40_.

### 2.4. Paralog pair estimation

The mRNA paralogous pairs for *Saccharomyces cerevisiae* are estimated based on the sequence similarities and identical domain content at the proteome level. First, we implemented NCBI-standalone version of protein-protein blast[50] across the proteomes. Blast hit pairs with E^−10^ *e-value* are further considered for filtering based on identical protein domain (*p < 10* ^*−5*^) assignment by Pfam[51](presently Interpro) (**Data S1**).

### 2.5 Statistical analysis

All the statistical tests provided in this manuscript are performed using PAST v3.0[52]. Generalized linear model (GLM) is performed using glm() function of RStudio.

## 3. Results and Discussion

### 3.1 Translationally efficient mRNA lives longer

During translation, aminoacyl-tRNA (aatRNA) is moved from the ribosomal A-site to the P-site and then to the E-site before being expelled (AA/PP/EE). This movement is asynchronous, as expulsion of uncharged tRNA from E-site is independent of incoming charged tRNA at the P-site [53]. Often times, slow decoding rate for mRNA due to abundance of NOC, increases the probability of attaining --/PP/-- state which attracts co-translational surveillance complexes like Ccr4-Not or Dhh1 [54,55] to initiate either COMD or NGD. The N-terminal domain of Not5 (component of Ccr4-Not), can sense the prolonged E-site vacancy triggering mRNA transcript release and degradation. Cryo-EM structure of a monosome fraction reveals Not5 binds to a vacant E-site in a conformation when there is no aatRNA at the A-site (--/PP/--) [56]. As a result of this functional overlap between NOC, COMD and NGD, we observe a positive correlation between genome-scale mRNA half-lives and translation rates (Pearson r = 0.55, *p =* E−158, **Fig. 1A**). Hence, it can be concluded that transcripts with higher translation rates can evade stalling of ribosomes, and thus exhibiting higher genome-level half-lives.

**Fig. 1.**
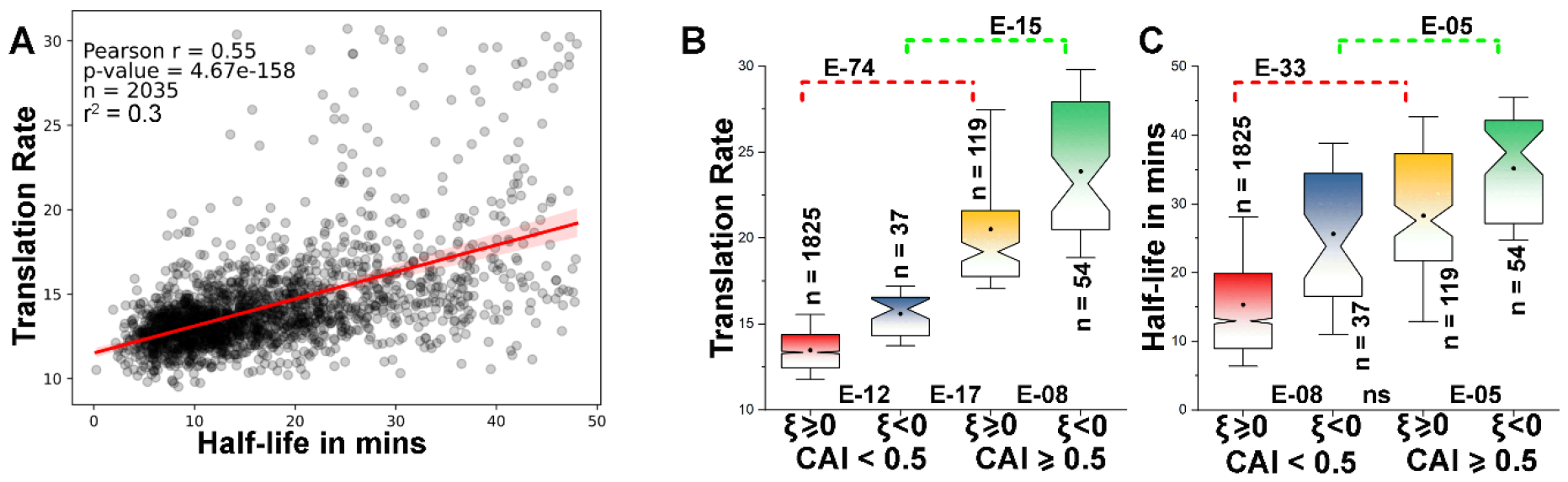
**(A)** Scatter plot showing the correlation between mRNA translation rates and half-lives. The red line states the best-fit line for the two sets of data. Notched box-plot showing distribution of **(B)** translation rates and **(C)** half-lives between structured (ξ<0) and unstructured (ξ ≥ 0) transcripts at low (CAI < 0.5) and high (CAI ≥ 0.5) CAI. The analysis for **(B)** and **(C)** are performed using ξ _s60us40_. The plot for ξ _s40us60_ is given in supplementary. Distributions in **(B)** and **(C)** are compared using Pairwise Mann-Whitney U-test and their respective *p*-values are provided.

### 3.2 Variations of mRNA translational rate and half-lives depend on both CAI and IUS

Since translation rates and half-lives exhibit strong associations among themselves, therefore, we look forward in understanding whether there is any trade-off between the parameters regulating these two processes. The initial choice of parameters are CAI[17] and IUS (ξ)[12], which are observed to be strong regulators of translation rates (Pearson r = 0.97, *p <* 10^−28^), and mRNA half-lives (Pearson r = − 0.59, *p <* 10^−28^) respectively. Owing to the interdependency among the two processes, we also observed a strong correlation between CAI with half-lives (Pearson r = 0.53, *p <* 10^−28^) and ξ with translation rates (Pearson r = − 0.61, *p <* 10^−28^) indicating that both mRNA translation rates and half-lives are regulated by more than one factors. The fact that a highly structured mRNA is efficiently translated could be due to the strong intrinsic helicase activity before translation, which can unwind stable structured CDS[57]. On the other hand, unstructured segments are prone to coil on itself, posing greater challenges for helicase activity and consequently diminishing both translation rates and half-lives. Recently, a study on designing efficient mRNA-based vaccine have elucidated that decreasing the number of U’s (uridine) in loops and hairpins within a transcript to generate a linear double stranded mRNA (ξ < 0 as per our calculation) can elevate mRNA stability and protein expression[58].

To gain a deeper insight, we grouped the transcripts as structured (ξ < 0) and unstructured (ξ ≥ 0), and each of these groups are subclassified based on low (CAI < 0.5) and high (CAI ≥ 0.5) optimality. We observed significant increase in both translation rates and half-lives of mRNA transcripts, with increasing structuredness (at fixed CAI), and with increasing CAI (at fixed ξ) (**Fig. 1B and C**), indicating the effect of both CAI and ξ on both the processes. We observed similar trend for translation rates and half-lives for ξ _s40us60_ (**Fig. S2A and B**).

Next, we have classified transcripts into six different groups based on the CAI values, and for each group, divided the transcripts into two different subgroups based on their structuredness. Since, the minimum and maximum CAI values for our set of transcripts are 0.16 and 0.91, we normalized the CAI score from 0 (minimum CAI) to 1 (maximum CAI). For, each of the classes, we have plotted the half-lives and translation rates for the mRNA transcripts having (i) ξ < M and (ii) ξ ≥ M (where M denotes median value) **(Fig. 2A and B)**. We observe that mRNA half-lives and translation rates gradually increase with CAI **(Fig. 2A and B)** for both ξ < M and ξ ≥ M. But, for a fixed CAI, transcript with higher internal un-structuredness (ξ ≥ M) exhibits lower stability and translation rates compared to transcript with lower internal un-structuredness (ξ < M). Unstructuredness greatly impact half-lives of mRNAs with lower optimality (Group 1-4 compared to Group 5 and 6), suggesting a dependency of endonuclease activity during NGD [9,41](**Fig. 2A**). On the other hand, at low range of CAI (group 1), the difference on structuredness and unstructuredness of the mRNA doesn’t affect the translation rates significantly (*p =* ns, **Fig. 2B**). With increase in CAI, the differences in translation rates between structured and unstructured transcripts, gradually increase (**Fig. 2B**).

**Fig. 2.**
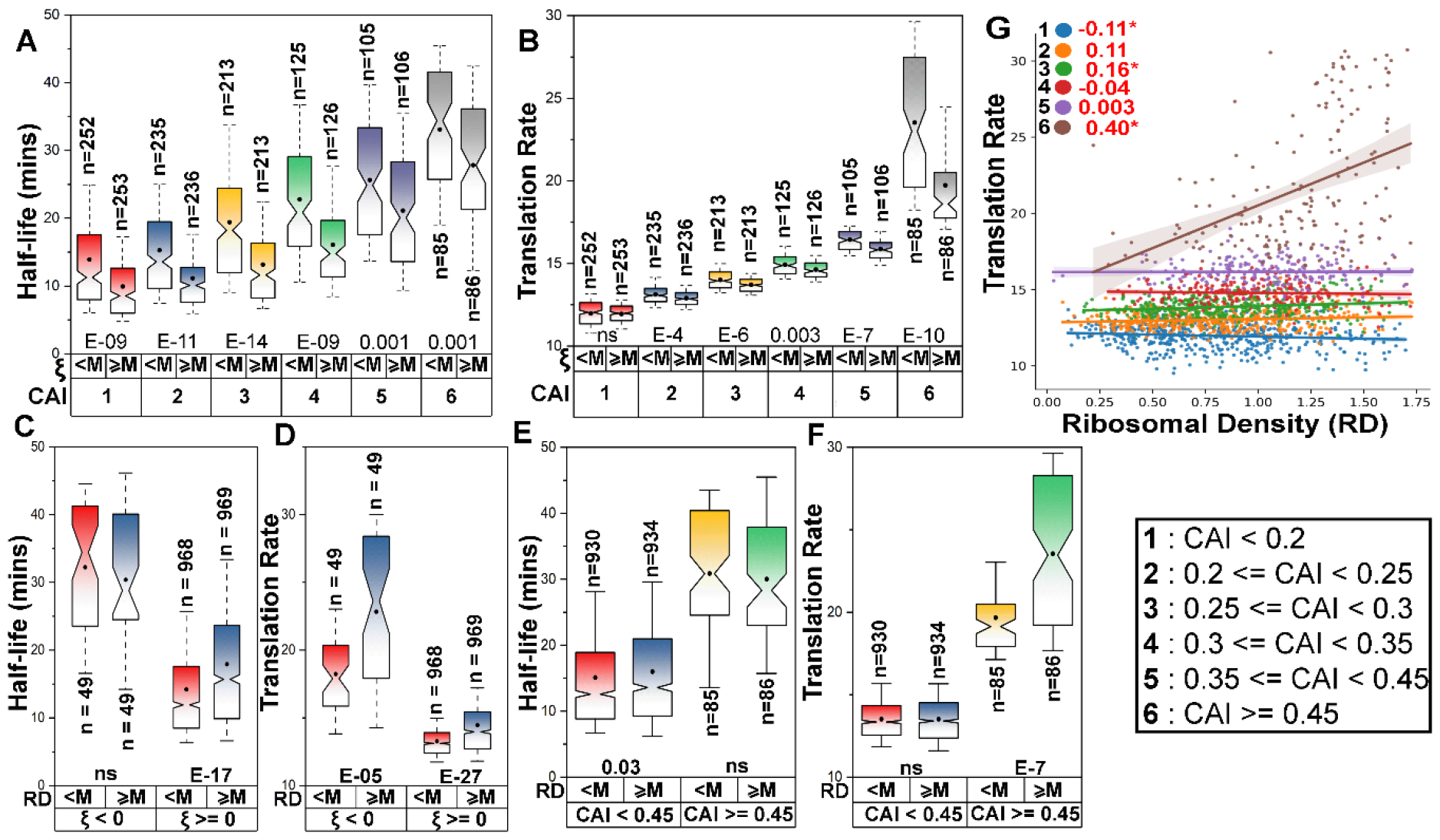
**(A and B)** The joint effect of CAI and ξ on **(A)** Half-lives and **(B)** Translation rates. At a fixed CAI range, the distribution of half-lives and translation rates is shown as notched box plot for different IUS range. For the ease of class separation, the CAI is normalized from 0 to 1 and is segregated into six classes. For each CAI class, the IUS values are partitioned based on the median (M) value of the ξ distribution. With increase in CAI, mRNA stability and translation rates increase. But within a same CAI group, structured transcripts exhibit a higher half-lives and translation rates compared to unstructured ones. Presence of unstructured segments facilitates endonuclease action and faster decay once the transcript is released from translational system. **(C and D)** The joint effect of IUS and RD on **(C)** half-lives and **(D)** translation rates. Here, the transcripts are segregated into structured (ξ < 0) and unstructured (ξ ≥ 0). For each ξ, the RD is divided into two classes based on their median. RD have no effect on half-lives of structured mRNA, while, it provides added stability to mRNA which are unstructured. For translation rate, with increase in RD, increases translation rates significantly at a fixed ξ range. **(E and F)** Effect of CAI and RD on **(E)** half-lives and **(F)** translation rates. Here we combined CAI classes 1-5 into one group, while the 6^th^ class represent the second group. For each group, we segregated mRNA based on their RD score (separated by median). We observe no significant change in mRNA half-lives when we combine CAI and RD. However, at low CAI, increase in RD doesn’t affect translation rate, while at higher CAI, increase in RD rapidly elevates the translation rates. Higher CAI indirectly indicates abundance of optimal codons, and upon higher ribosomal density, largely increases the translation rates. **(G)** Scatter plot showing the correlation among RD and translation rates for different CAI classes. Pairwise Mann-Whitney test is done for **A, B, C, D, E and F** and the *p*-values are provided below the notched boxes. All the analysis are performed using ξ _s60us40_ and Ribozero (RD dataset).

We have already observed decrease in both mRNA translation rates and half-lives with increase in unstructuredness (ξ). But within a fixed ξ range, transcripts with higher optimal codons (CAI ≥ M) exhibit higher translation rates and half-lives compared to those with lower optimal codons (CAI < M). Moreover, the impact of higher CAI is more prominent for the unstructured transcripts (**Fig. S3A and B**).

The fact that CAI and ξ look highly intertwined among themselves in regulating both mRNA translation rates and half-lives is due to the various endonucleases which are responsible to trigger cleavages of a larger mRNA fragments into smaller by-products upstream of the stall regions[59] during NGD or COMD. It has been reported that the stability of NGD-cleaved mRNA is largely dependent on endonucleases like Dom34/Hbs1 complex [41,42,60]. Once cleaved, the 5′-end products can be degraded by 5′-exonucleases, while the 3′ -ends can be degraded by exosomes[41].

### 3.3 The context specific effect of RD on mRNA translation rates and half-lives

#### 3.3.1 RD increases translation rates and half-lives but to a certain limit

Ribosomal density (RD) is a measurement of the “translational fitness” of a given mRNA in response to various other environmental and developmental factors[61]. Generally, mRNA loaded with heaviest polysomes are thought to have highest translational efficiency and lower stability[58]. This indicates a presence of an optimal point beyond which the synchronized movement of the polysomes is hampered [62], leading to overcrowding and collision between the ribosomes resulting in translational dependent mRNA decay[63]. To understand the impact of increasing RD on mRNA translation rates and half-lives, our comprehensive genome scale analysis, reveals a gradual elevation of both parameters initially (**Fig. S3D and E**). However, beyond a certain threshold, while translation rates continue to rise marginally, mRNA half-lives start to decline (**Fig. S3D and E**).

#### 3.3.2 RD can only harbor stability to unstructured mRNA

A further study showed that ribosomes can only impart stability to mRNA transcripts which are unstructured (ξ ≥ 0) (Mann Whitney (MW) *p =* E − 17, **Fig. 2C**). No significant difference is observed for structured mRNA (ξ < 0) with increase in ribosomal density (**Fig. 2C)**. This is further assisted by correlation analysis indicating no significant relation between RD and half-lives on structured mRNA transcripts (Pearson r = − 0.003, *p =* ns, **Fig. S3F**), whereas a very weak correlation (Pearson r= 0.2, *p =* E − 19, **Fig. S3F**) on unstructured transcripts.

#### 3.3.3 RD can elevate translation rates of both structured and unstructured transcripts

On the other hand, RD can elevate translation rates for both structured and unstructured transcript (**Fig. 2D**). The median translation rates for structured transcripts (ξ < 0) are significantly higher than unstructured ones (ξ ≥ 0) indicating that the ribosomal translocations are more efficient and streamlined in structured transcripts compared to the unstructured ones [64,65]. In agreement with the last observation, the pearson correlation between RD and translation rates is higher for the structured transcripts (Pearson r = 0.48, *p =* E − 07, **Fig. S3G**) than unstructured ones (Pearson r = 0.3, *p =* E − 42, **Fig. S3G**). And within a same ξ- group, transcripts harboring more ribosomes have a higher translation rates (**Fig. 2D**).

#### 3.3.4 Effect of RD on translation rates is observed at higher CAI

Next, we tried to observe the collective effect of CAI and RD on mRNA half-lives and translation rates (**Fig. 2E and F**). We observe no significant difference for mRNA stability with increasing RD at both low and high CAI ranges (*p =* ns, **Fig. 2E)**. However, at low CAI, although increased RD doesn’t elevate TR, but a significant change is observed at higher CAI (MW *p =* E − 07, **Fig. 2F**). A lower CAI is indicative of abundance of NOC within a transcript, which has a longer wait period owing to their lower concentration of cognate aa-tRNA. We found that transcripts with the higher CAI (CAI > 0.45) exhibits a high positive correlation (Pearson r = 0.4, *p =* E − 08) between RD and translation rates (**Fig. 2G**) compared to other CAI classes.

Frequency of occurrence of optimal codons (CAI) within a mRNA transcript reflects its intrinsic translational efficiency within the codon level. The results indicate that if intrinsic translational efficiency is lower (lower CAI), then increasing in RD can’t elevate translation rates. On the other hand, higher RD can elevate translation rates of transcripts enriched with optimal codons (higher CAI).

### 3.4 Partial correlation analysis reveals CAI and IUS as major determinants for translation rates and half-lives respectively

The indication of optimal codons, unstructured segments and RD regulating both mRNA translation rates and half-lives motivates us to look for their relative contributions. Here, we first estimated pearson correlations to understand the linear causality. However, when multiple confounding parameters regulate a process, a linear correlation doesn’t always confirm causality. To overcome this limitation, we also implemented a parallel partial correlation analysis.

It is evident from **Fig. 3A** that CAI has a significant pearson correlation with both translation rates (r = 0.97, *p =* 0) and mRNA half-lives (r = 0.53, *p =* E − 150). However, when we performed the partial correlation analysis, the correlation coefficient for CAI with translation rates drops by 25% (Partial r = 0.73, *p =* 0), while it almost vanishes for mRNA half-lives (Partial r = − 0.06, *p =* 0.005). Similarly, for ξ (**Fig. 3A**), we also observed a strong negative correlation with mRNA translation rates (Pearson r = − 0.61, *p =* E − 206) and half-lives (Pearson r = − 0.59, *p =* E− 190). However, this high negative correlation for ξ against mRNA translation rates completely vanishes in partial correlation (r = −0.06, *p =* 0.01), and we observe a 40% decrease from their pearson r to their respective partial r (r = −0.33, *p =* E−51). In addition to CAI and ξ, we also included tAI, tCSC and RD in this analysis (**Table S1**). The drops in partial correlation and their comparison suggest that ξ and CAI are the major regulators of mRNA half-lives and translation rates respectively.

**Fig. 3.**
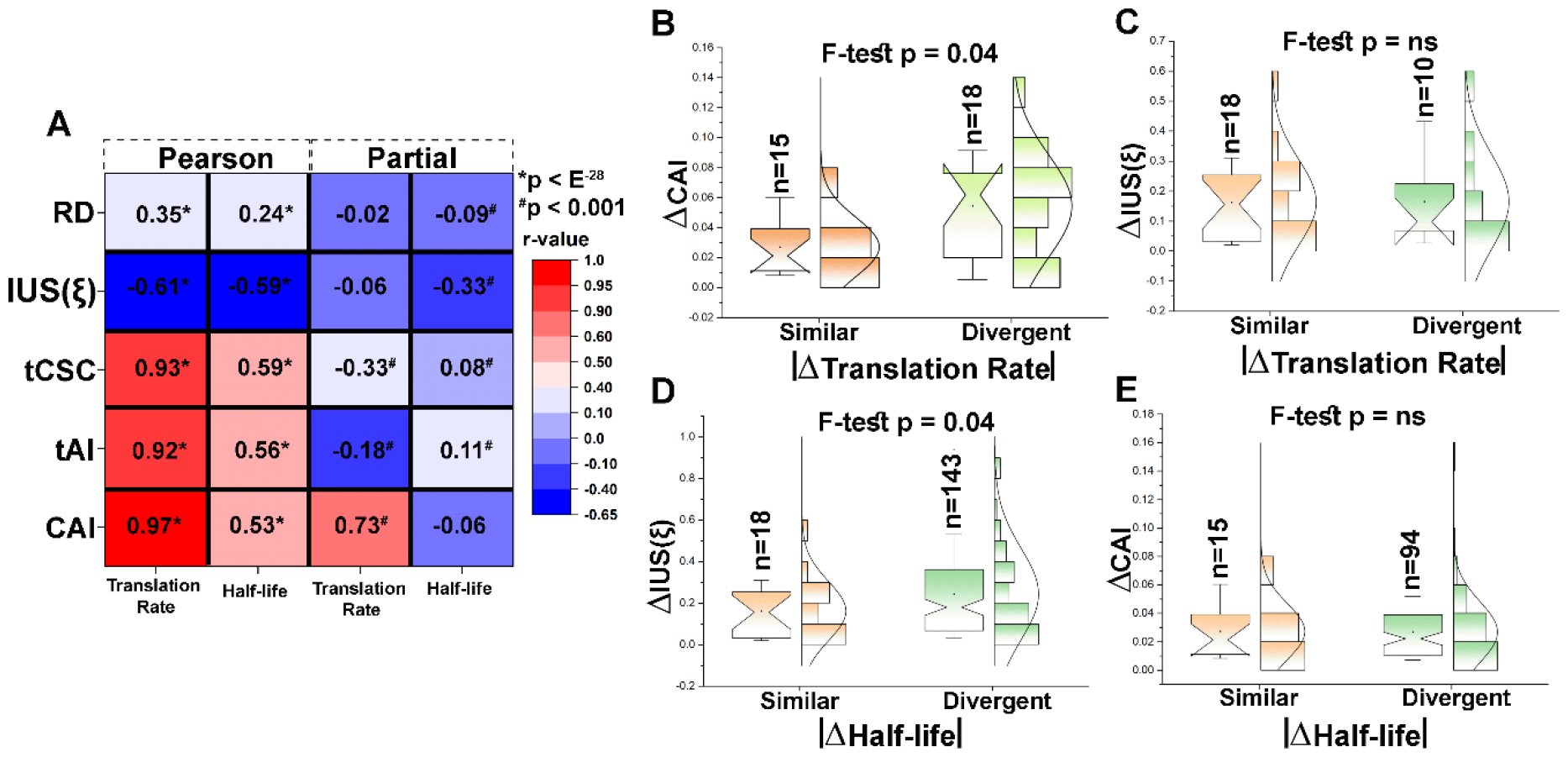
**(A)** Heatmap showing the linear correlations (using Pearson r) and partial correlations of all the studied sequence (CAI, tAI, and tCSC), structural parameters (ξ) and RD (Ribozero) with translation rates and half-lives. The ‘*’ and ‘^#^’ used indicates the significance level for the pearson and partial correlation respectively. Partial correlation analysis indicates CAI and IUS standout as the primary regulator of mRNA translation rates and half-lives respectively, among all the others. The complete table with the exact *p*-values is provided as Table S1 in supplementary. Distribution plot of **(B)** ΔCAI at similar ξ and half-lives and **(C)** Δξ at similar CAI and half-lives for paralogous pairs having similar and divergent translation rates. Distribution plot of **(D)** Δξ at similar CAI and translation rates and **(E)** ΔCAI at similar ξ and translation rates for paralogous pairs having similar and divergent half-lives. From **(B), (C), (D) and (E)**, it is observed that CAI is important for change in translation rates and ξ is important for regulation of half-lives. All the analysis performed here are done using ξs60us40, while the analysis for ξs40us60 is provided as Fig. S4E, F, G and H in supplementary. The similar and divergent distributions are tested using F-test and their respective *p*-values are provided.

We further constructed a generalized linear model (GLM) to quantify the relationship between (i) Translation rates and (ii) Half-lives with their predictors namely RD, tCSC, TAI, IUS and CAI. For translation rate, CAI is observed to be the best predictor (*p =* 0, t-statistics) compared to others (**Table S2A**). In case of half-lives, IUS is observed to be the best predictor (*p =* E−55, t-statistics) (**Table S2B**).

### 3.5 Paralogous transcripts with differential translation rates and half-lives show differences in CAI and IUS respectively

Gene duplication often led to rise of new copies featuring altered structural geometries. Previously, we had reported that much of the half-lives’ variation among the paralogous proteins and the mRNA encoding the paralogous proteins are an outcome of the altered structural geometry and oligomerization criteria [12,66]. Therefore, we hypothesized that the altered CAI and ξ among paralogous transcripts would lead to altered translation rates and mRNA half-lives respectively.

To test our hypothesis, we initially calculated the absolute difference for CAI, IUS, translation rates and half-lives for the paralogous pairs. We designated paralogous pairs as (i) similar, if the transcript pairs fall within a strict range of CAI and IUS (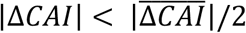 or 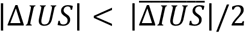) and (ii) divergent, if the transcript pairs show wide variation of CAI and IUS 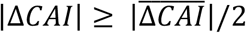 or 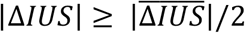. For the complete set of paralogous pairs (641 pairs), we found significant differences in distribution of Δ(translation rate), among similar and divergent CAI and ξ pairs (F-test pCAI = E−62 and F-test pξ = E−36). The distribution for the divergent pair is much wider compared to the similar ones (**Fig. S4A**). On the other hand, in case of Δ(half-lives) we observed significant change in the distributions for similar and divergent ξ (F-test *p =* E−12, **Fig. S4B**), but CAI is deemed to be non-significant. Similar pattern is observed for similar and divergent Δξ _s40us60_for both translation rates and mRNA half-lives (**Fig. S4C and D**).

It should be noted that the paralogous pairs may have differences in both CAI and ξ, which influence the mRNA translation rates and half-lives either individually or in conjunction. Therefore, we further intensified our selection criteria for performing the same analysis. We assigned the paralogous pairs as similar and divergent based on their differences in translation rates and half-lives and compared the distributions of CAI (for similar ξ, **Fig. 3B** and **3E**) and ξ (for similar CAI, **Fig. 3C** and **3D**). It is evident from **Fig. 3B** and **3C**, that the difference in translation rates among the paralogous pairs (at similar half-lives) is mainly regulated by the difference in CAI. Similarly, the divergence in half-lives within the paralogous pair having similar translation rates are primarily due to the variation in internal unstructured segments (**Fig. 3D and E**). This analysis confirms that CAI and ξ are the major determinants for the divergence on translation rates and half-lives respectively along the evolutionary time scale post gene-duplication. This pattern remains consistent, when reanalyzed with Δξ _s40us60_ (**Fig. S4E, F, G and H**)

## 4. A complex interplay between codon optimality and mRNA structuredness regulate mRNA translation rates and half-lives

Our results illustrate that, despite mRNA translation and degradation being distinct cellular processes, their regulations remain closely linked through a common set of factors including occurrence of optimal codon, proportional of internal unstructured segments and ribosomal density. Among all the factors, codon optimality is the major determining factor governing translation rate, but half-lives are primarily controlled by internal unstructured segments. Additionally, analysis of paralogous pairs further confirms these major determinants (half-lives = *f* (ξ) and Translation rates = *f*(CAI)) at the evolutionary scale.

Co-translational surveillance complexes like Ccr4-Not/Dhh1 are closely linked to rare codons at the A-site of ribosomes, particularly in slow decoding rates[56]. This triggers the recruitment of mRNA decay factors, including various (endo)-exonucleases, emphasizing the connection between lower mRNA translation rates and early degradation. Here comes the role of unstructuredness of the translation stalled mRNAs into their degradation. Translation-stalled mRNAs with higher unstructuredness are more susceptible to endonuclease[12] binding and early degradation, resulting in shorter half-lives. Previously it was observed that in *S. cerevisiae*, an endonuclease, Cue2, targets mRNA around the A-site of stalled ribosomes, further making the cleaved transcripts available for degradation by 3’ and 5’- exonucleases [9,14,67]. Thus, our analysis highlights the interplay of codon optimality and mRNA structural characteristics in influencing translationally coupled mRNA decay/COMD/NGD, showcasing a synchronized regulatory mechanism.

Since mRNAs are inherently unstable, improving the stability is a primary objective for increasing the effectiveness of an mRNA vaccine[68]. Identification of factors for regulating mRNA translation rates and stability would provide a recommender system to design mRNA with maximum potency[69]. With the recent paradigm shift towards mRNA-based vaccine, these foundational principles could be later used to create mRNA-based drug delivery system for development of prophylactics and therapeutics studies which can be actively being investigated against a broad spectrum of disease.

## 8. Authors Contribution

SB, SH and SK designed the research and implemented the computational methodologies. SB performed the research and analyzed the data. SB and SH wrote the initial draft. SB, SH and SK reviewed and edited the manuscript.

