## Supplementary for "Translation coupled mRNA-decay is a function of both structural and codon level characteristics"

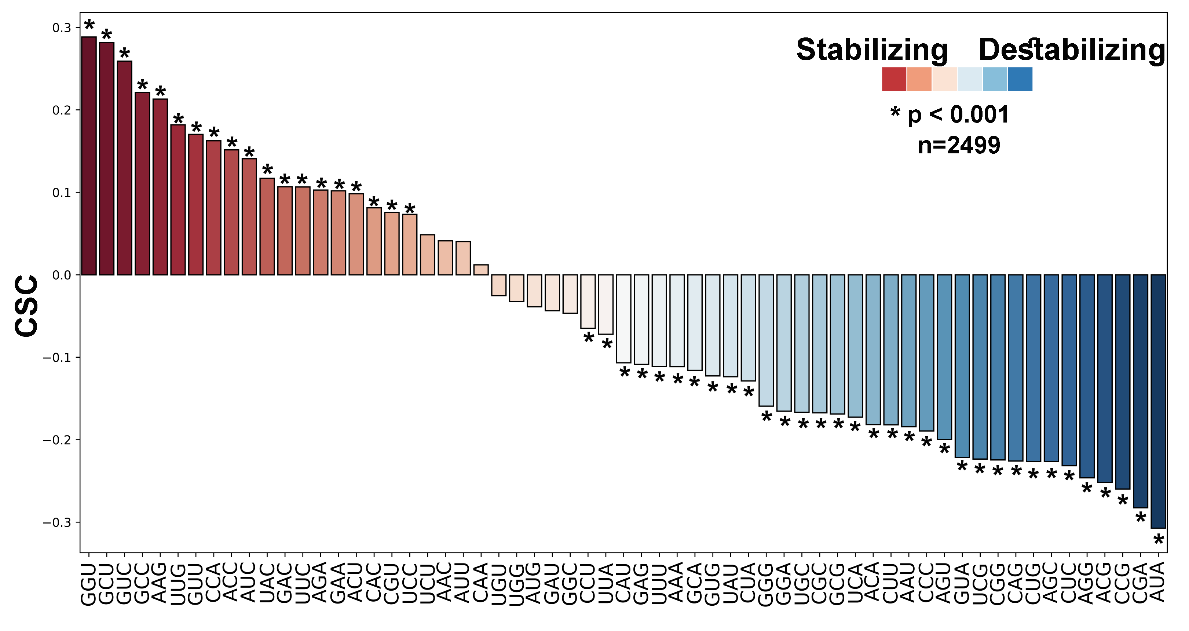

Fig. S1A. Codon stabilization coefficient (CSC) plot for all the 61 codons for the yeast transcriptome. CSC is computed as the pearson correlation coefficient between the frequency of each codon within a transcript with their respective half-lives. The codons which are deemed to significant (*p* < 0.001) are marked as ‘*’.

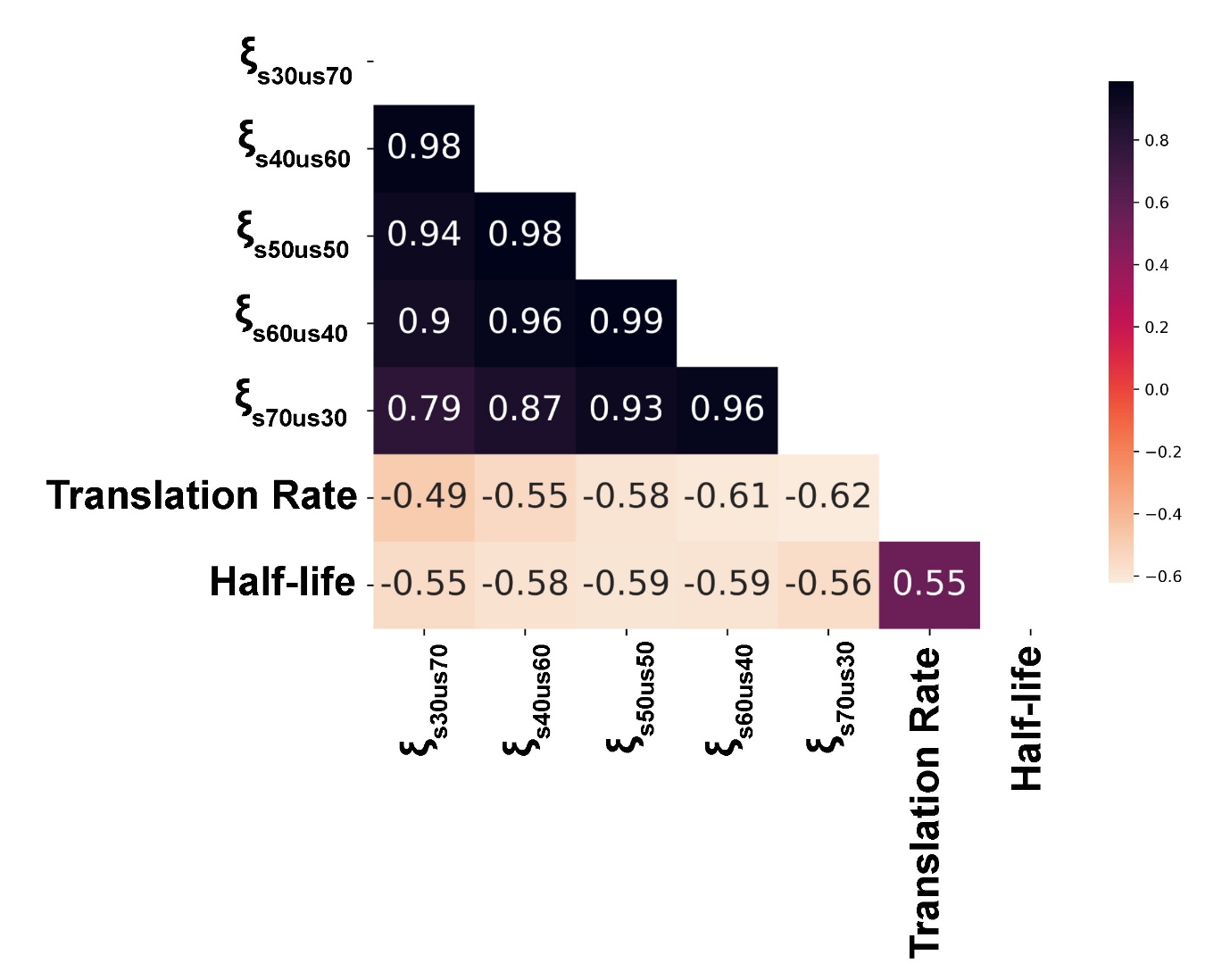

Fig. S1B. Pearson correlation matrix showing the collinearity between ξ calculated at different structured and unstructured threshold for a window. The matrix also shows the pearson correlation of ξ (at different structured and unstructured threshold) with translation rates and mRNA half-lives. Since ξ (at different threshold) are highly correlated among themselves and with translation rates and mRNA half-lives, hence, we used a representative ξ_s60us40_ for further analysis. We have also generated the results using ξ_s40us60_, showing that the pattern does not changes.

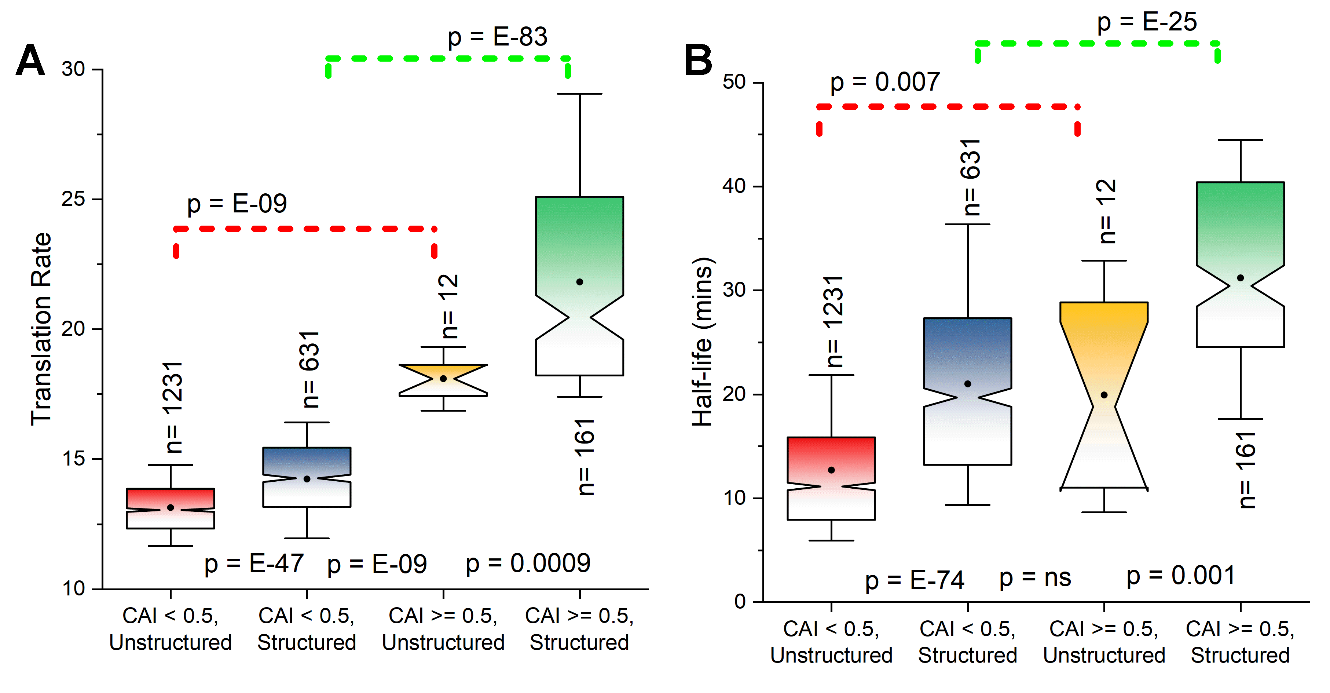

Fig. S2. Notched box-plot showing distribution of **(A)** translation rates and **(B)** half-lives between structured (ξ_s40us60_<0) and unstructured (ξ _s40us60_ ≥ 0) transcripts at low (CAI < 0.5) and high (CAI ≥ 0.5) CAI. Distributions are tested for pairwise Mann-Whitney U-test for medians and their respective *p*-values are provided. The plot for ξ _s60us40_ is given in Fig. 1B of main text. The analysis for the Coller half-life dataset [Presnyak et. al., 2015] for ξ _s60us40_ is given in Data S1.

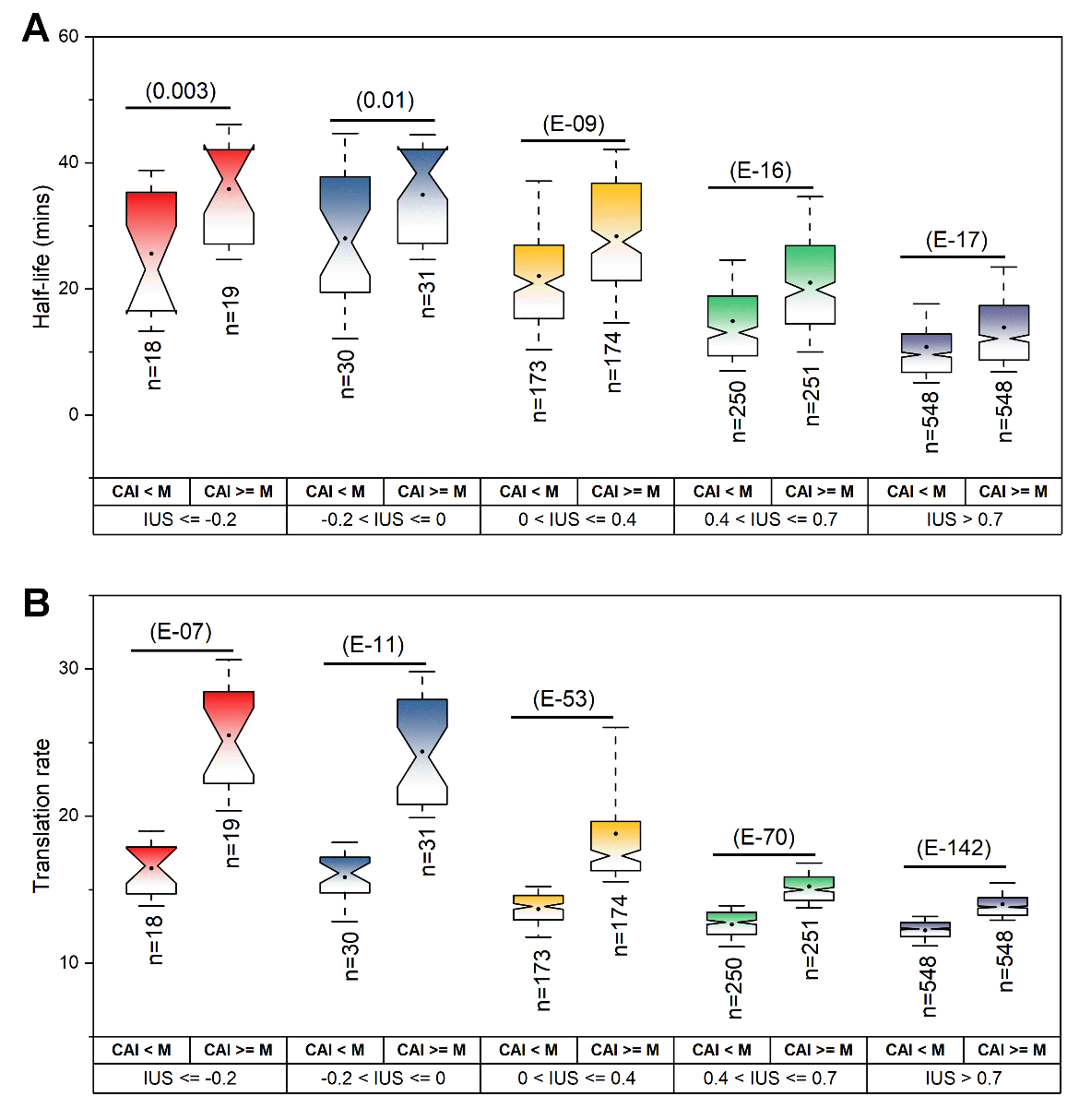

Fig. S3A, B. Effect of increasing CAI at a fixed range of internal unstructuredness (ξ) on mRNA (A) half-lives and (B) translation rates. We observe decrease in mRNA translation rates and half-lives with increasing unstructuredness of mRNA. Within each IUS group, the effect of CAI on mRNA half-lives and translation rates are much more prominent for unstructured mRNA compared to structured ones. Distributions within a group are tested with pairwise Mann-Whitney U-test and their *p*-values are provided. Here, the analysis is done with ξ_s60us40_. The analysis for ξ_s40us60_ for both translation rates and half-live (Cramer data) [Miller et. al., 2011] is given in Data S1. The analysis using the Coller dataset [Presnyak et. al., 2015] for ξ_s60us40_ is given in Data S1.

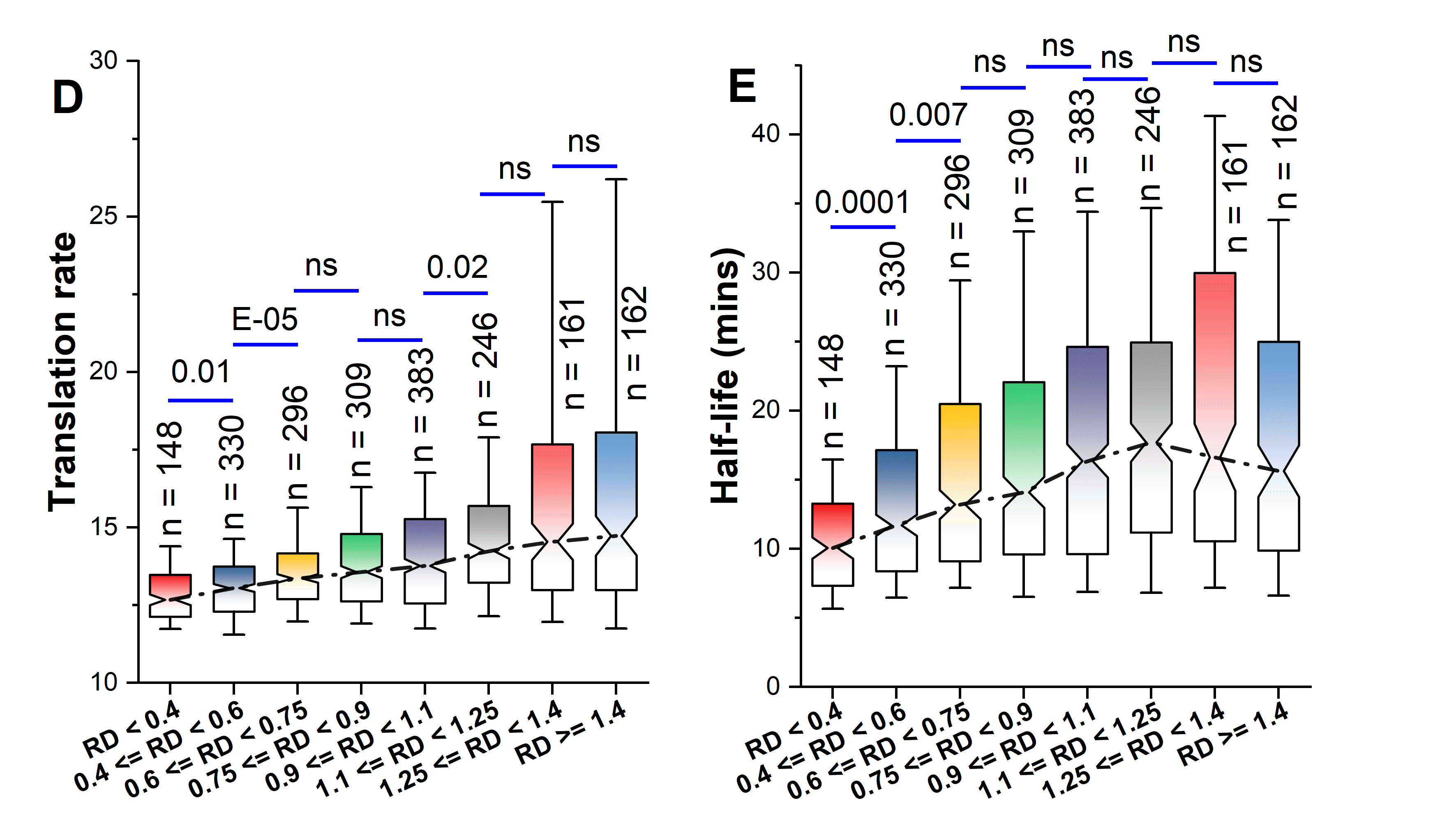

Fig. S3D and E. Effect of increasing RD on (D) mRNA translation rates and (E) half-lives. Increasing RD initially increases the translation rates significantly upto a certain limit. On the other hand, increasing RD increases the half-lives initially, but beyond a certain limit, half-lives decrease gradually.

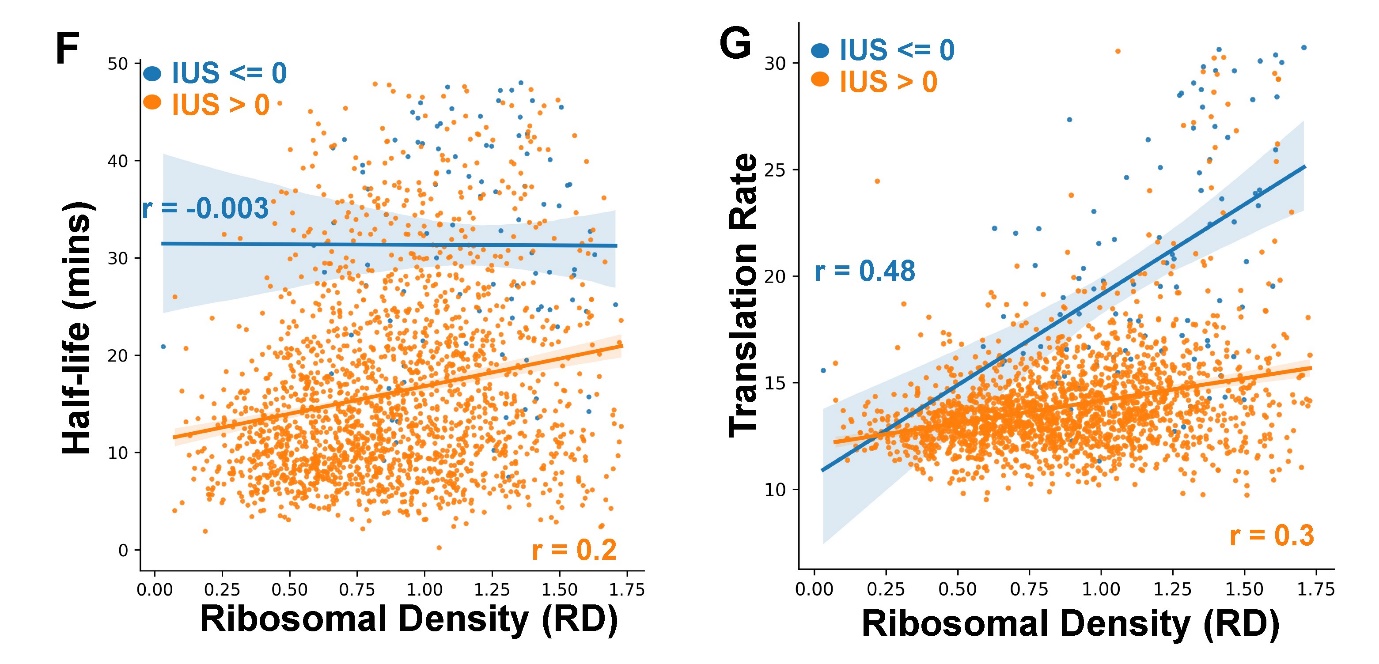

Fig. S3F and G. Scatter plot indicating the change in mRNA (F) half-lives and (G) translation rates with increasing RD for both structured and unstructured transcripts. The pearson correlation values (r-value) are provided for both structured (blue) and unstructured (orange) transcripts. Here, ξ_s60us40_ is used for this analysis.

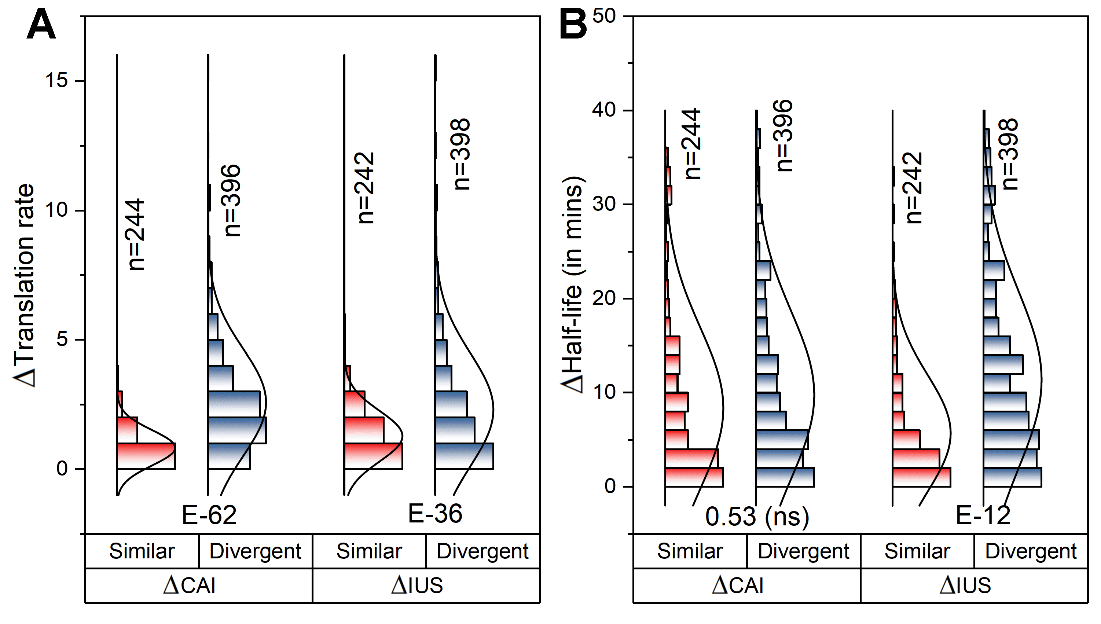

Fig. S4A and B. Paralogous pairs are classified into two categories Similar and Divergent (Detailed description for Similar and Divergent criteria given in main text). Distribution plot showing the difference in (A) translation rates and (B) mRNA half-lives for paralogous pairs having similar and divergent CAI and IUS. For (A), the difference in translation rates is a result of the difference of both CAI and IUS, while for (B), the difference in half-lives is a result of only IUS. In case of half-lives, CAI is deemed to be non-significant. The distributions are tested with F-test and their respective *p*-values are provided. Here, the analysis is done using ξ_s60us40_.

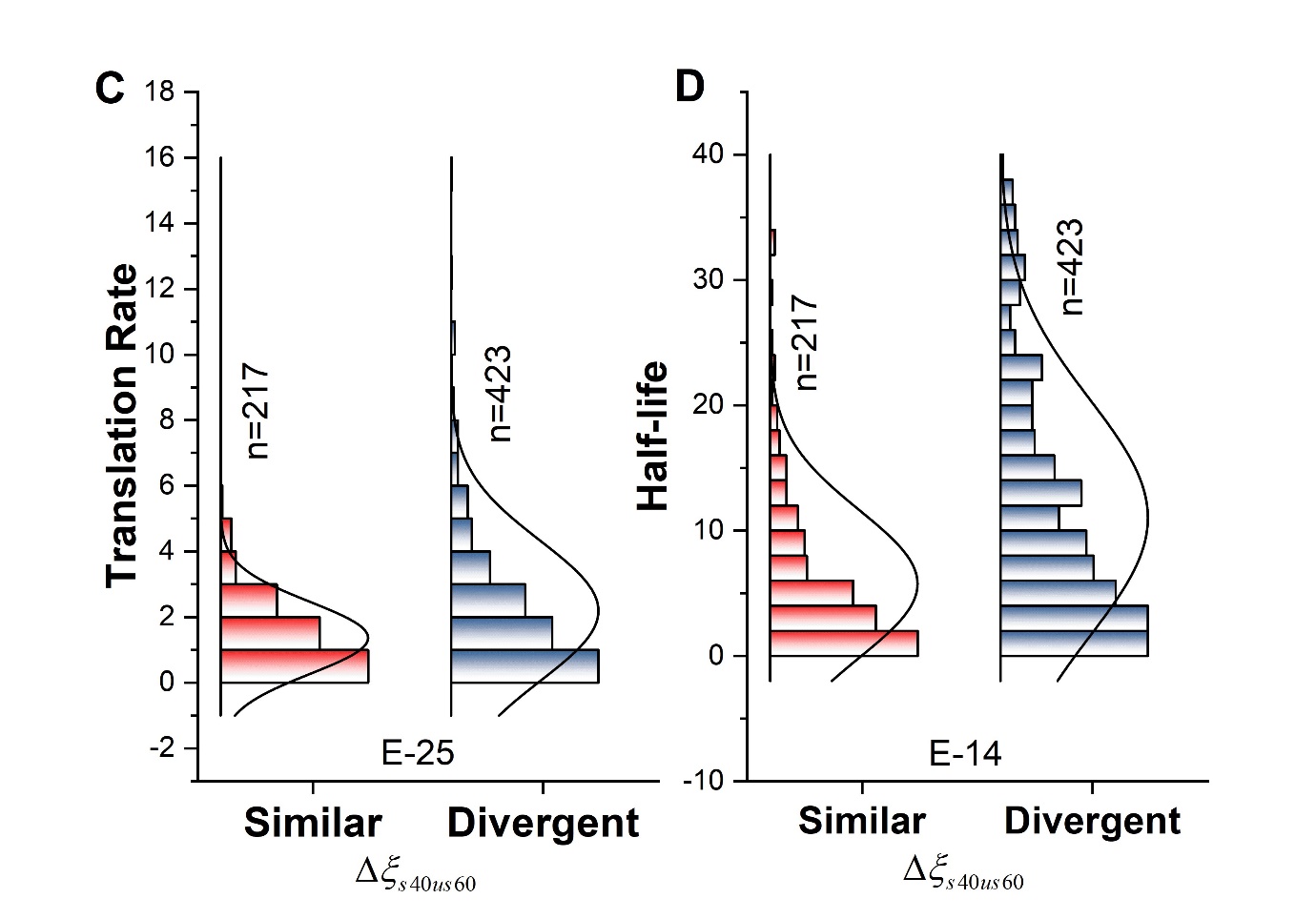
Fig. S4C and D. Paralogous pairs are classified into two categories Similar and Divergent (Detailed description for Similar and Divergent criteria given in main text). Here, distribution plot showing the difference in (A) translation rates and (B) mRNA half-lives for paralogous pairs having similar and divergent ξ_s40us60_. The plot for ξ_s60us40_ is given in Fig. S4A and B. The distributions are tested with F-test and their respective *p*-values are provided.

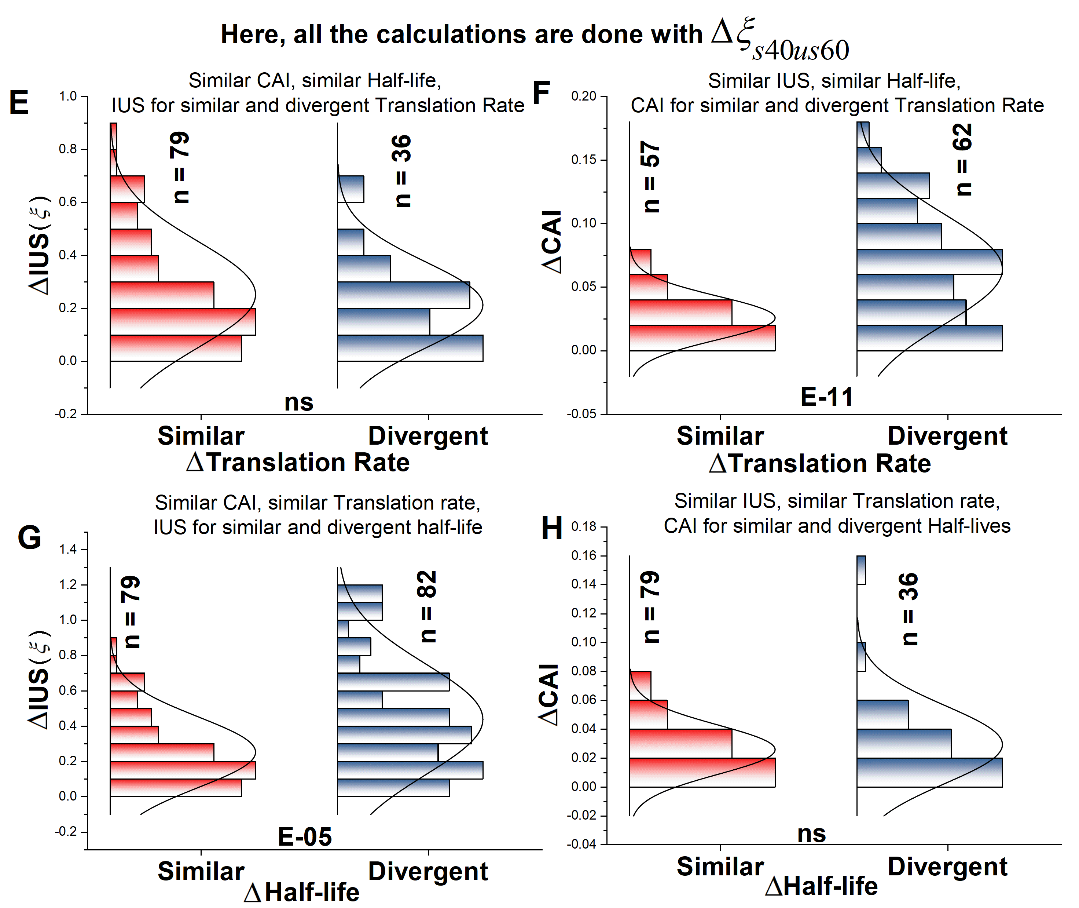

Fig. S4E, F, G and H. Distribution plot of **(E)** ΔIUS (ξ) at similar CAI and half-lives and **(F)** ΔCAI at similar IUS(ξ) and half-lives for paralogous pairs having similar and divergent translation rates. Distribution plot of **(G)** ΔIUS (ξ) at similar CAI and translation rates and **(H)** ΔCAI at similar IUS(ξ) and translation rates for paralogous pairs having similar and divergent half-lives. From **(E), (F), (G) and (H)**, it is observed that CAI is important for change in translation rates and IUS(ξ) is important for regulation of half-lives. All the analysis performed here are done using ξ_s40us60_. The analysis for ξ_s60us40_. The similar and divergent distributions are tested using F-test and their respective *p*-values are provided.

|  | **Pearson** | | | | **Partial** | | | |
| --- | --- | --- | --- | --- | --- | --- | --- | --- |
|  | **Translation Rate** | ***p*-value** | **Half-life** | ***p*-value** | **Translation Rate** | ***p*-value** | **Half-life** | ***p*-value** |
| **CAI** | 0.96594 | 0 | 0.53329 | 6.21E-150 | 0.72945 | 0 | -0.06107 | 0.005918 |
| **tAI** | 0.91606 | 0 | 0.56143 | 2.23E-169 | -0.18281 | 1.03E-16 | 0.10922 | 8.12E-07 |
| **tCSC Length** | 0.92702 | 0 | 0.59398 | 2.43E-194 | 0.33121 | 3.59E-53 | 0.075662 | 0.000645 |
| **IUS (ξ)** | -0.60823 | 3.39E-206 | -0.58948 | 1.02E-190 | -0.055481 | 0.012416 | -0.32673 | 1.04E-51 |
| **RD** | 0.3541 | 3.566E-61 | 0.24066 | 3.346E-28 | -0.022977 | 0.3008 | -0.08667 | 9.24E-05 |

Table S1. Complete table of Pearson and Partial correlation along with their respective *p*-values for CAI, tAI, tCSC, IUS and RD with mRNA translation rates and half-lives. All the codon dependent parameters tCSC, tAI and CAI show very high pearson correlation with translation rates. The comparative drop in the partial correlation values from their respective pearson correlation in all these three cases suggest CAI act as a best codon dependent descriptor for translation rates. IUS show moderates negative pearson correlation with translation rate, but diminishes in the partial correlation. IUS, tCSC, tAI and CAI shows a good pearson correlation with mRNA half-lives. While IUS is observed to show 40% decline from pearson to partial, tCSC, CAI and tAI almost vanishes for half-life. Thus, IUS is a major indicator for half-lives. RD shows a weak pearson correlation with both translation rates and half-lives, which diminishes in partial correlation.

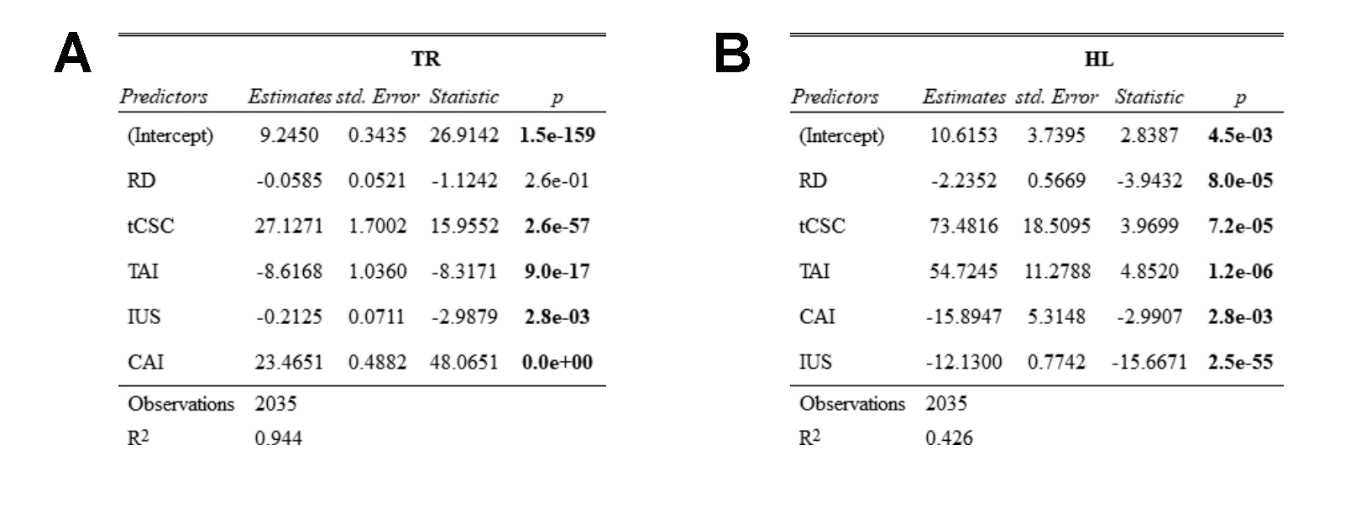

**Table S2**. Generalized Linear Model (GLM) analysis performed for (A) Translation rate (TR, dependent variable) and (B) Half-life (HL, Dependent variable) with their predictors (independent variables namely RD, tCSC, TAI, IUS and CAI) to understand their relative importance. The “(Intercept)” indicates the null model, which does not contain any predictors. The number in “Estimates” indicates the change in the dependent variable with unit change in the predictors. For example in (A), for RD, ̶ 0.06 change in TR is associated with one unit change in RD. The “std. Error” indicates the error associated with the Estimates. Here, “Statistic” indicates the t-statistics which is the ratio between Estimates and std. Error. Therefore, a higher t-statistics along with the associated “*p*” indicates a better predictor. Here for (A) we observed that CAI is the best determinant for regulation of translation rates (t-statistics *p* = 0), and (B) IUS has the best determinant for regulation of half-lives (t-statistics *p* = 2.5E ̶ 55). This result is generated using glm() function of RStudio.
